# Gait-symmetry-based human-in-the-loop optimization for unilateral transtibial amputees with robotic prostheses

**DOI:** 10.1101/2020.10.04.325720

**Authors:** Yanggan Feng, Chengqiang Mao, Qining Wang

## Abstract

Gait asymmetry due to the loss of unilateral limb increases the risk of injury or progressive joint degeneration. The development of wearable robotic devices paves a way to improve gait symmetry of unilateral amputees. Moreover, the state-of-the-art studies on human-in-the-loop optimization strategies through decreasing the metabolic cost as the optimization task, have met several challenges, e.g. too long period of optimization and the optimization feasibility for unilateral amputees who have the deficit of gait symmetry. Here, in this paper, we proposed gait-symmetry-based human-in-the-loop optimization method to decrease the risk of injury or progressive joint degeneration for unilateral transtibial amputees. The experimental results (*N* = 3 unilateral transtibial subjects) demonstrate that only average 9.0±4.1*min* of convergence was taken. Compared to gait symmetry while wearing prosthetics, after optimization, the gait symmetry indicator value of the subjects wearing the robotic prostheses was improved by 21.0% and meanwhile the net metabolic energy consumption value was reduced by 9.2%. Also, this paper explores the rationality of gait indicators and what kind of gait indicators are the optimization target. These results suggest that gait-symmetry-based human-in-the-loop strategy could pave a practical way to improve gait symmetry by accompanying the reduction of metabolic cost, and thus to decrease the risk of joint injury for the unilateral amputees.

## 1. Introduction

Human-in-the-loop approaches are the state-of-the-art optimization method, which are inspired from humans to optimize walking performance unintentionally Alexander (1989); Selinger and Donelan (2014). Most existing human-in-the-loop approaches in the fields of wearable robots improve the walking performance through reducing the metabolic cost Ding et al. (2018); Zhang et al. (2017). For example, Zhang *et al*. proposed a human-in-the-loop optimization method for a bilateral ankle exoskeleton that reduced the average 24.2% metabolic cost Zhang et al. (2017). Then the torque pattern could be automatically tuned and customizing individual parameters become possible. Ding *et al*. implemented a human-in-the-loop optimization of hip assistance with a bilateral soft exosuit during walking Ding et al. (2018). Through optimizing the peak and offset timing, an average of 21.4*min* and metabolic cost reduction by 17.4% were found, compared with walking without the device. The above human-in-the-loop application mainly focused on the wearable exoskeleton, which often optimized the metabolic cost as the main task. Human-in-the-loop approaches exhibit the potential ability for the personalized wearable robots in a wider application Walsh (2018). However, requiring lengthy evaluation periods Ding et al. (2018); Selinger and Donelan (2014); Zhang et al. (2017), time-varying dynamics of human behaviors Selinger et al. (2015) and complicated neural and cognitive development Makin et al. (2017), raised the challenge of human-in-the-loop objective functions based on the reducing the metabolic cost rate. Moreover, few studies of human-in-the-loop focused on the unilateral amputees, who had the consequences of gait symmetry.

Biological studies show that due to the loss of lower limb, the amputees exhibited the gait asymmetry in kinetics and kinematics, compared to the able-bodied Sanderson and Martin (1997). For the unilateral amputees, this kind of gait asymmetry influences the maximum torque and torque steadiness significantly in the intact leg in some extents Simoneau-Buessinger et al. (2019). The reason for gait asymmetry maybe mainly resulted from strength asymmetry Lloyd et al. (2010), during walking. Strength asymmetry in unilateral transtibial amputees has a moderate relationship in an increased risk of developing osteoarthritis Lloyd et al. (2010). For the transtibial amputees, the walking speed also influenced gait symmetry Nolan et al. (2003). During jumping landing, peak force asymmetry and moment asymmetry may potentially be easy to result in injury or progressive joint degeneration for transtibial amputees and improving gait symmetry in jumping is beneficial to avoid excessive loading onto one or the other limb Melzer et al. (2001); Schoeman et al. (2013). Gait asymmetry also contributes to function disability, and therefore improving gait symmetry to avoid the potential risk of illness is a challenge Patterson et al. (2010). Also, many gait indicators in terms of kinetics and kinematics are proposed to evaluate the gait symmetry of transtibial users or asymmetry Kendell et al. (2010). However, efforts to reduce kinematic asymmetries have not resulted in improved symmetries in the underlying kinetics Mattes et al. (2000); Childers and Kogler (2014), and thus selecting a suitable gait symmetry also becomes a challenge. Unilateral amputees due to the reduction of afferent signal from proprioreceptors, may result in deterioration in walking stability Viton et al. (2000) and malfunction of selected muscles Combs et al. (2013). Walking instability due the limb loss may lead to a fall, even musculoskeletal injury Svoboda et al. (2012). Further, improving kinetic symmetry could overcome walking instability in some extent Kendell et al. (2010); Svoboda et al. (2012).

Therefore, targeting on unilateral lower-extremity amputees, this paper investigated the kinetic-gait-symmetry-based human-in-the-loop online optimization. the main contribution of this paper was as follows. Firstly, this optimization method mainly focused on the unilateral amputees and implement on-line prosthesis parameter optimization. Moreover, reducing optimization duration and thus customizing the individual parameters also are studied. In addition, this paper focuses on metabolic cost during optimization and studies the consistency relationship between metabolic cost and gait symmetry.

## 2. Diagram of Gait-Symmetry-Based Human-in-loop optimization

Human-in-the-loop optimization based on reducing metabolic cost in the fields of wearable robots made some progress Ding et al. (2018); Felt et al. (2015); Zhang et al. (2017). Subjects in these studies are mainly from healthy subjects and all of these subjects have a good performance on gait symmetry between both limbs. However, for the unilateral amputees, gait asymmetry will increase the risk of injury or illness Lloyd et al. (2010), and thereby improving gait asymmetry will be also necessary for these kinds of subjects. Meanwhile, reducing the optimization periods and thereby improving optimization efficiency is worthy to explore Ding et al. (2018); Selinger and Donelan (2014); Zhang et al. (2017). Therefore, targeting on the unilateral amputees, this paper demonstrates one human-in-the-loop-based gait symmetry optimization.

The process of gait-symmetry-based human-in-the-loop optimization is as shown in Fig. 1. First, three unilateral amputees wearing robotic prostheses were asked to walk on the treadmill at 1.0*m/s*. The treadmill could measure ground reaction forces (GRFs) of both feet while walking. Next, the gait symmetry indicator was calculated using GRFs. Further, the Bayesian optimization algorithm was utilized to search the optimal control parameters. Finally, the optimal control parameters were transferred to the robotic prostheses via Bluetooth and used to control prostheses. The process would continue until the optimization converged. Repeat the above process until the optimization process converged.

**Figure 1.**
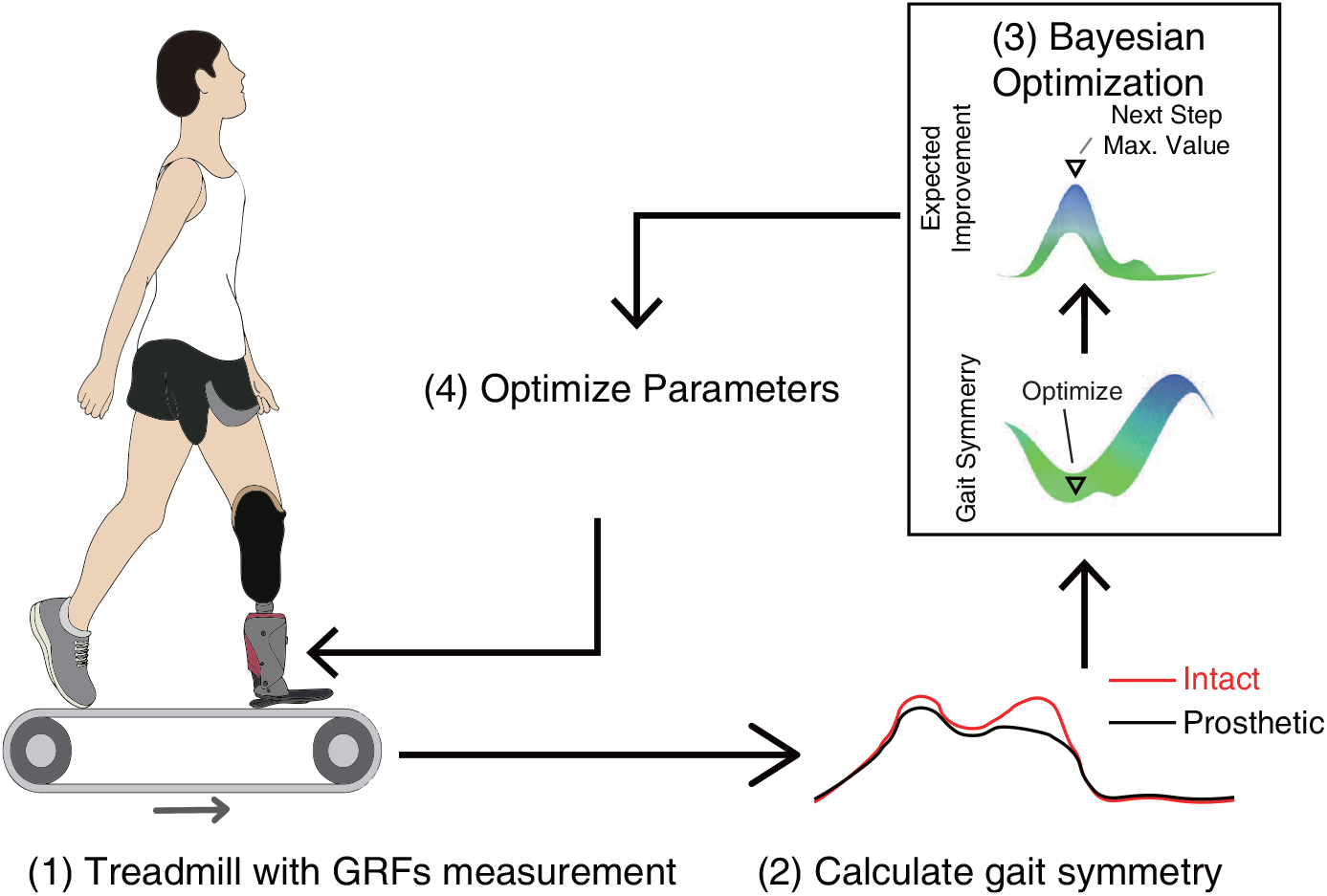
Optimization process. First, the subject wearing a robotic prosthesis is asked to walk on the treadmill at 1.0*m/s*. The treadmill can measure GRFs while walking. Second, based on GRFs, the gait symmetry indicator is calculated. Third, Bayesian optimization is utilized to obtain the next control parameters. Fourth, the control parameters are updated.

## 3. Robotic Prosthesis

### 3.1 Prosthesis prototype

A light-weight prosthesis PKU-RoboTPro-Plus was used, which is an extended version from our previous studies Wang et al. (2015); Feng and Wang (2019). The electrical control system comprised the sensor subsystem, the microcontroller subsystem and the actuator subsystem. The sensor subsystem was composed of a strain gauge bridge sensor, an angle sensor, a motor encoder and a switcher (details can refer to Feng and Wang (2019); Feng et al. (2019)). A strain gauge bridge sensor was utilized to measure the interaction force between the carbon-fiber footplate and ground during stance phase. An angle sensor, a motor encoder and a switcher were used for ankle joint measurement, motor speed measurement and power supply on/off switching, respectively. The microcontroller unit utilized STM32F103 processor (ARM Cortex M3) to deal with sensor data and control the actuator. The actuator subsystem utilized a three-phase motor driver (A4931, Allegro LLC.) to implement both passive braking torque control and active position control. A 50 *W* motor (EC 45, Maxon Inc.) was utilized to actuate the prosthesis. A laptop communicated with the robotic prosthesis via Bluetooth and the communication baud rate was set 57600 *bit/s*.

### 3.2 Control strategy

Based on the gait phase (stance phase and swing phase), the motor controller was designed to implement different control strategies (braking torque control and position control).

- Braking Torque Control. We proposed a method to implement realtime prosthetic braking torque control Wang et al. (2015). The methods can result in the equal braking torque *τ*_*eb*_ due to the different duty cycle (*D*_*p*_) of the Pulse Width Modulation (PWM) signal and the equation can be estimated as follows:

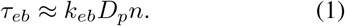

where *k*_*eb*_ is the proportionality coefficient in the unit of *Nm/rpm* and *n* is the motor speed in the unit of *rpm*. As the duty cycle *D*_*p*_ determines the relationship between the torque and the speed, it can be regarded as the damping coefficient.
- Position Control. For the active motion state, the Proportional-Derivative (PD) controller was developed in motor control. The principle of the PD controller is as follows:

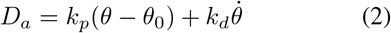

where *D*_*a*_ is the active controller output for motor driver, *k*_*p*_ and *k*_*d*_ are predefined constants, *θ*_0_ is the desired equilibrium angle, *i.e*. ankle angle when heel strike. In this paper, *k*_*p*_ and *k*_*d*_ are set to 6 and 2 respectively.

From Eq. 1, we can see that parameters *D*_*p*_ has an effect on the ankle braking torque of robotic prosthesis. Eq. 2 demonstrates that *θ*_0_ affects the prosthetic kinematics. Parameters *D*_*p*_ and *θ*_0_ have a certain range of value, in Fig. 2. The determination of the control parameters range depends on the individual’s preferences. More detailed description about control strategies could refer to our previous study Feng and Wang (2019); Feng et al. (2019); Wang et al. (2015).

**Figure 2.**
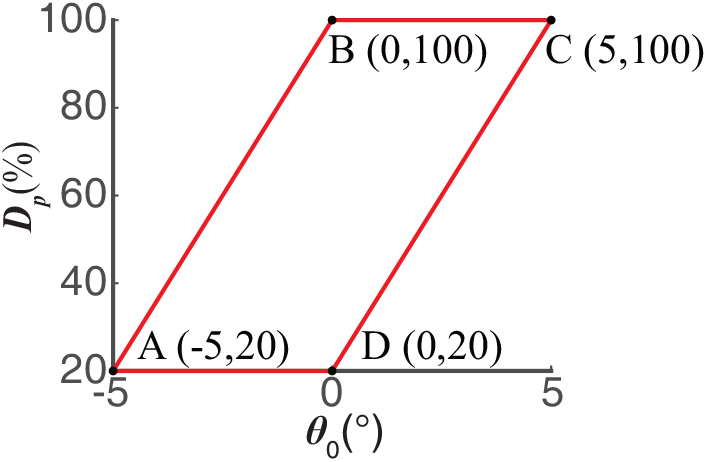
The range of control parameters. When the shank is perpendicular to the plane of the carbon-fiber footplate, the ankle angle is defined as 0°. The positive angle represents the dorsiflexion and the negative angle represents the plantar flexion. A, B, C and D stand for four boundary condition points, respectively.

## 4. Gait Symmetry Calculation

### 4.1 Definition

The gait symmetry indicator paves a way to evaluate the walking performance of subjects. The gait symmetry indicator used in our experiments is defined as the root mean square value of the GRFs difference between the left and right foot in a gait cycle as follows,

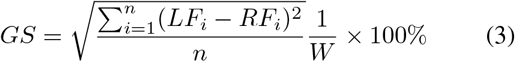

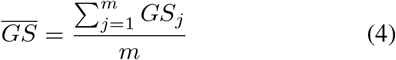

where *GS* represents the gait symmetry indicator for each stride, 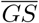 represents the average gait symmetry indicator of *m* strides, *LF*_*i*_ represents the GRFs of the left foot at the *i*-th sampling time point in a gait cycle, and *RF*_*i*_ represents the GRFs of the right foot at the *i*-th sampling time point in a gait cycle, and *W* represents the gravity of the subject to normalized, and *n* represents the total number of sampling points of the GRFs of left and right feet in one gait cycle. When the values of *LF*_*i*_ and *RF*_*i*_ are closer to the same, it indicates that the gait symmetry of the unilateral transtibial amputees while wearing the robotic prosthesis is better, and the *GS* is also smaller.

### 4.2 Transition time

Considering that the active prosthesis switches between different control parameters, the unilateral transtibial amputees also need to spend a certain transition time to adapt to the new control parameters of the robotic prosthesis. Therefore, we first conducted a preliminary experiment to measure the transition time. As shown in Fig. 3, in the 120th and 240th gait cycles (red dotted line), a new set of control parameters is sent, and it can be observed that the transition time requires an average of 15 gait cycles. Then, the average value of the gait symmetry indicator of the 10 gait cycles (*m* = 10 in Eq. 4) after the transition time is taken as the gait symmetry indicator 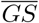 corresponding to the control parameter.

**Figure 3.**
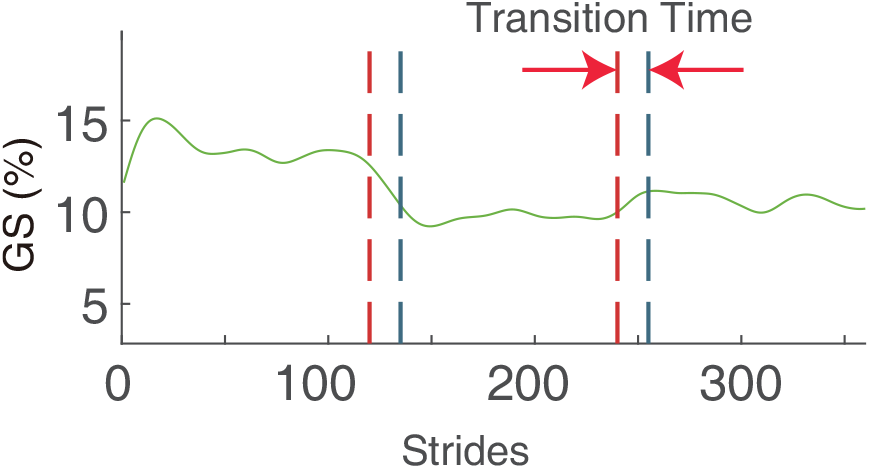
Strides represent the number of gait cycles, GS represents the gait symmetry indicator, the red dotted line indicates that a new set of control parameters was sent at the time, and the blue dotted line indicates that new control parameters begin to function at that time.

## 5. Bayesian Optimization

Bayesian optimization is a global optimization method, especially suitable for the case where the correspondence between the objective function and the optimization parameters is not well written into the analytical form Brochu et al. (2010); Mockus et al. (1978). It first assumes that the prior distribution of the objective function is in accordance with the Gaussian distribution and then obtains new data according to the iterative process. According to more information, the posterior distribution of the objective function can be obtained, so that the posterior distribution of the objective function approach the real objective function. The Bayesian optimization algorithm has a good balance between exploration and exploitation when searching between the feasible domains of the optimization parameters Brochu et al. (2010).

Bayesian optimization algorithm can be formalized as follows:

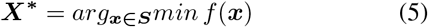

In the Eq. 5, ***S*** is a candidate set of ***x***. The goal is to choose an ***x*** from ***S*** such that the value of *f* (***x***) is the smallest. It is possible that the specific formula of *f* (***x***) cannot be known, but if an ***x*** is selected, the value of *f* (***x***) can be obtained.

In our experiments, ***x*** = [*D*_*p*_, *θ*_0_], where *D*_*p*_ can be regarded as the damping coefficient, and *θ*_0_ represents the desired equilibrium angle. ***S*** represents the feasible domain of the control parameter ***x***, that is, the range of values of *D*_*p*_ and *θ*_0_. *f* (***x***) represents the gait symmetry indicator 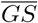 under the control parameter ***x*** = [*D*_*p*_, *θ*_0_]. The optimization algorithm of our experiment is divided into the following four steps.

### 5.1 Parameter initialization

We first divide the feasible domain ***S*** of the *D*_*p*_ and *θ*_0_ into six regions, and then select a set of control parameters ***x*** = [*D*_*p*_, *θ*_0_] from each region pseudo-randomly, then get six sets of initial control parameters ***x***_**1**_, ***x***_**2**_, …, ***x***_**6**_, and then under the six sets of control parameters, the experiment is performed to obtain the corresponding gait symmetry indicator *f* (***x***_**1**_), *f* (***x***_**2**_), …, *f* (***x***_**6**_). We want to ensure that the control parameters between the two regions are not very close. When we divide the six regions, there is still a certain interval between the regions and the regions, as shown in Fig. 4.

**Figure 4.**
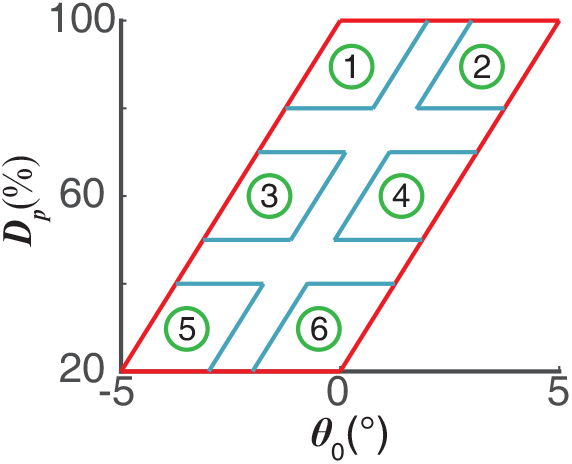
The feasible domain of the control parameters is evenly divided into six parts, and the initial points are randomly selected from the six regions.

### 5.2 Gaussian process regression

The prior distribution of the gait symmetry indicator *f* (***x***) satisfies the Gaussian process, that is, *f* (***x***) ∼ *GP* (***µ*(*x*), *k*(*x, x*,))**, where ***µ*(*x*)** represents the expected value ***E*(*f* (*x*))** of the sample *f* (***x***), which is usually taken as zero. ***k*(*x, x***^***′***^) is a covariance function, usually with an anisotropic square exponential kernel function as a covariance function Brochu et al. (2010)

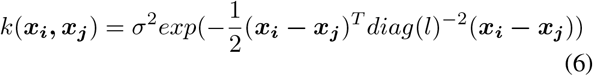

where *σ*^2^ is the gait symmetry indicator 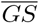 variance and *l* is a diagonal matrix consisting of the length scale parameters of *D*_*p*_ and 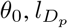 and 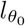. 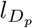and 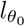 measure the sensitivity of the gait symmetry indicator in the *D*_*p*_ and *θ*_0_ dimensions, respectively.

Assume there is now a set of initialized sample points ***D***_**1:*t***_ **= { *x***_**1:*t***_, ***f***_**1:*t***_ **}**, and for a new sample *x*_*t*+1_, the joint Gaussian distribution is:

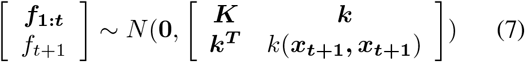

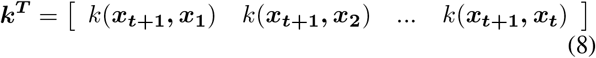

where ***K*** is the positive definite kernel matrix **[*K***_***ij***_] = *k*(***x***_***i***_, ***x***_***j***_), (*i, j* = 1, 2, …, *t*). Considering the noisy case, the distribution of noise is considered to be 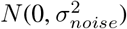, We can get the posterior gait symmetry indicator distribution of *f*_*t*+1_:

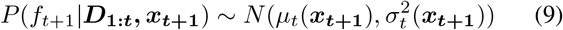

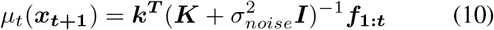

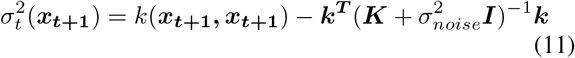

We use the maximum log-likelihood method to obtain the hyperparameter 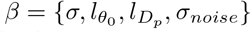 value by selecting six sets of initial values.

### 5.3 Expected improvent function

After computing the gait symmetry indicator posterior indicator distribution given all data, the next *x* are selected by maximizing expected improvement (EI). the utility function *u*(***x***) as follows:

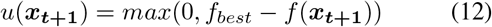

where *f*_*best*_ = *minf*_*i*_, *i* = 1, 2, …, *t*. The expected improvement function *EI*(***x***) as follows:

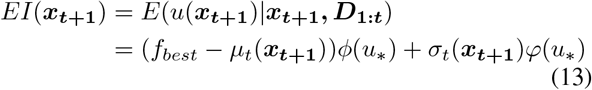

where *u*_∗_ = (*f*_*best*_ − *µ*_*t*_(***x***_***t*+1**_))*/σ*_*t*_(***x***_***t*+1**_), *f*_*best*_ = *minf*_*i*_, *ϕ*(*x*) is the cumulative distribution function of the standard normal distribution, *φ*(*x*) is the probability density function of the standard normal distribution Brochu et al. (2010). If *σ*_*t*_(***x***_***t*+1**_) = 0, *EI*(***x***_***t*+1**_) = 0. The next control parameter 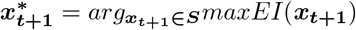. In general, the maximum problem can be converted to the minimum problem. In our experiments we used the fmincon function in MATLAB to find the minimum value, which might be the local minimum.

### 5.4 Convergence Condition

There are two convergence conditions (a) and (b). When one of the convergence conditions is satisfied, the optimization process ends.

a. If the gait symmetry indicator value *f* (***x***_***t***_) appearing at time *t* is the minimum or sub-minimum of all current values, and |*f* (***x***_***t***_)− *f* (***x***_***t*−1**_) |or |*f* (***x***_***t***_) −*f* (***x***_***t*−2**_) |is less than 10% of the maximum gait symmetry indicator value in the initial six values. it is considered that time *t* converges and the optimization process ends.
b. If the number of iterations exceeds 30, the optimization process also ends.

## 6. Optimization Experiment Procedure

### 6.1 Overview

The whole experiment consisted of four steps. First step-adaptation, the subjects wearing a robotic prosthesis were walked on the treadmill at 1*m/s* for at least 30 minutes, making the subject familiar with the prosthesis. Second step-initialization, the subjects wearing the robotic prosthesis, in given the initial six sets of control parameters, walked on the treadmill at 1*m/s* to complete the initialization process. Third step-post-iteration, based on the initialized result, the control parameters of the robotic prosthesis were optimized, and gait symmetry indicator *GS* was calculated simultaneously until completing optimization. Four step-metabolic cost evaluation, three sets of control parameters from the optimization process, were selected and the corresponding metabolic cost was recorded to evaluate the walking performance. The three sets of control parameters corresponding to the best gait symmetry (min condition), intermediate one (mid condition) and worst one (max condition) in the optimization process. During each condition, the subjects are asked to stand on the treadmill for at least 2 minutes and walk on the treadmill for at least 3 minutes at 1*m/s* in the three conditions(i: min condition, ii: mid condition, iii: max condition). In contrast to the walking performance of passive prosthesis, the subjects were asked to wear the passive prosthesis to walk in the same condition as the comparison (iiii: passive condition-wearing passive prostheses). For four conditions, a respiratory measurement device is used to measure the metabolic cost rate, and the optical capture system is used to measure the ankle joint angle and the knee joint angle of the subject.

### 6.2 Subject

Three subjects volunteered to participate in the experiment and wrote the informed consent before experiments. All experiments in this paper were approved by the local ethics committee of Peking University. Three subjects were also told the possible consequences in the process of the experiment in advance. They were blind to know the control parameters of the robotic prosthesis. The detailed information of three subjects was listed as Tab. 1.

**Table 1.** Detailed information about the amputee subjects.

| Subject | Age (years) | Weight (kg) | Height (cm) | Leg | Gender | Prosthesis | Prosthesis Weight (kg) |
| --- | --- | --- | --- | --- | --- | --- | --- |
| 1 | 38 | 65 | 170 | Left | Male | Otto Bock 1C60 | 0.460 |
| 2 | 57 | 81 | 170 | Left | Male | Otto Bock 1C30 | 0.345 |
| 3 | 31 | 72 | 171 | Right | Male | Otto Bock 1C40 | 0.420 |
| Mean | 39.2 | 69.8 | 170.8 |  |  |  | 0.408 |
| SD | 14.6 | 8.3 | 1.1 |  |  |  | 0.058 |

### 6.3 Communication Protocol between Treadmill and Prosthesis

Communication between the treadmill and the computer and communication between the computer and the robotic prosthesis is very important. Their normal communication with each other ensures the normal collection of the GRFs of left and right feet and the normal control of the robotic prosthesis. The treadmill, slave laptop and master laptop are first connected to the same router via LAN cable, so that data can be transmitted under the same LAN. The bottom of the treadmill is equipped with four six-dimensional force sensors. The treadmill can measure the GRFs of left and right feet of the subject during walking on the treadmill, and then transmit the data from the treadmill to the slave laptop. In the slave laptop, the mechanical data is read by installing special software (Gaitway-3D software). The master laptop establishes a connection with the slave laptop through the TCP/IP protocol, reads the GRFs of left and right feet of the subject during walking from the slave laptop. The master laptop calculates the gait symmetry indicator according to the data, and then utilizes MATLAB realizes Bayesian optimization algorithm. By completing the optimization process, it will get the optimized control parameters of the robot prosthesis. In the next step, the master laptop will send the optimized control parameters to the robot prosthesis through Bluetooth to realize the control of the robot prosthesis.

### 6.4 Wear and Measurement

Fig. 6 demonstrates that one subject wears robotic prostheses, a portable respiratory measurement unit and some markers to record motion trajectory walks on the treadmill. The angle between two extension lines of thigh and shank was defined as a knee joint angle after removing the initial drift. The angle between two extension lines of shank and foot was defined as an ankle joint angle after removing the initial drift. The angle two legs stood upright on the treadmill was an initial angle. A wireless portable respiratory measurement unit transferred the data of metabolic cost to a PC at each breath wireless. Simultaneously, the GRFs were record through two laptops and a transit router.

**Figure 5.**
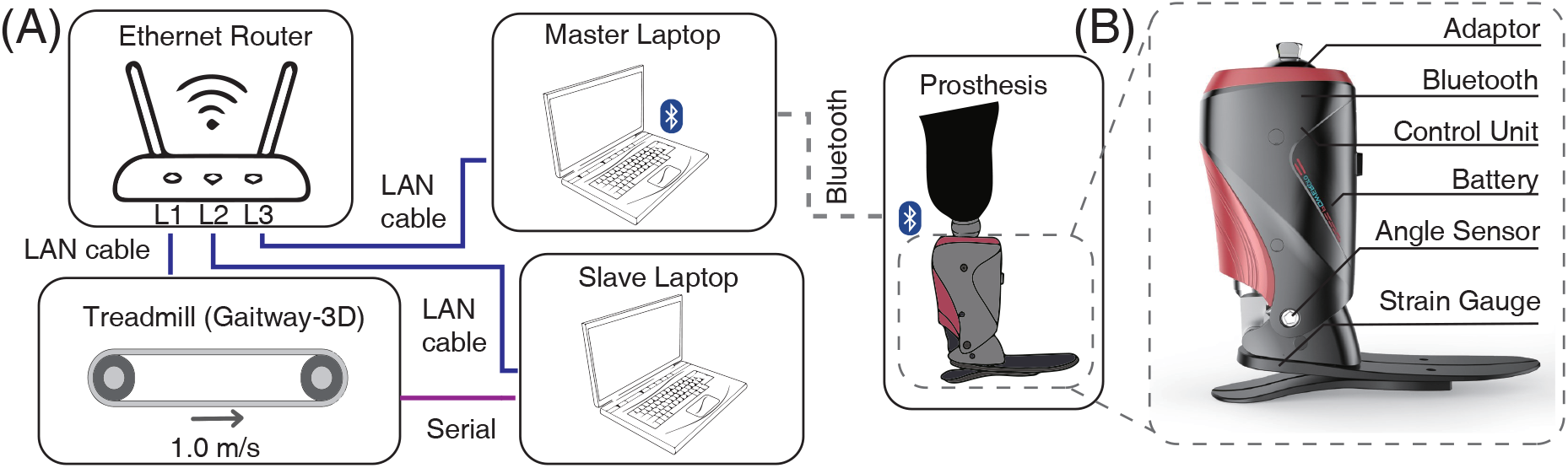
Communication protocol between setups and prosthetic prototype. (A) Communication protocol between setups. The slave laptop is connected to the treadmill via a LAN cable and utilized to collect GRFs on both feet using a professional software (Gaitway-3D software). The sampling frequency is 100*Hz*. Simultaneously, the slave laptop commands the treadmill speed via a serial cable. The master laptop obtains GRFs on both feet from an Ethernet router via a LAN cable. And then, the master laptop calculates gait symmetry and executes a Bayesian optimization using Matlab. After getting control parameter next step, the master will send it to the robotic prosthesis via a Bluetooth device. The LAN cable and serial cable and Bluetooth exchange data via the TCP/IP and RS232 protocol, respectively. (B) Prothetic prototype. The adaptor is a connector between the socket and prosthesis. Two types of sensors are utilized-an angle sensor to measure ankle joint angle and a strain gauge to detect gait events. The robotic prosthesis weighs 1.58*kg* (including the plugged-in rechargeable Li-ion battery). A Bluetooth device receives the commands from the master laptop and sends it to the control unit.

**Figure 6.**
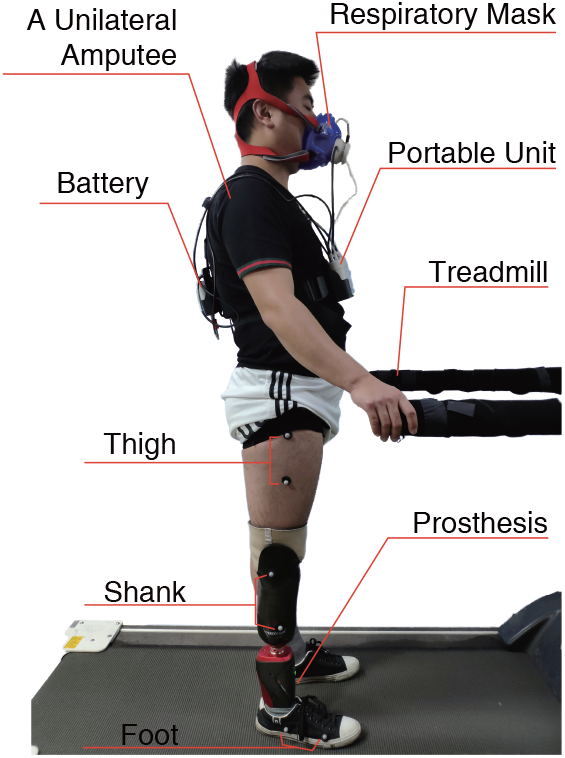
The subject’s wearing diagram and the physical instrument connection diagram.

## 7. Result

### 7.1 Convergence

For all subjects, the convergence time of the optimization experiment was 9.0±4.1*min*, and the convergence time included the experimental time of the six sets of initialization parameters and the time of the subsequent optimization process experiments, as shown in Fig 7.

**Figure 7.**
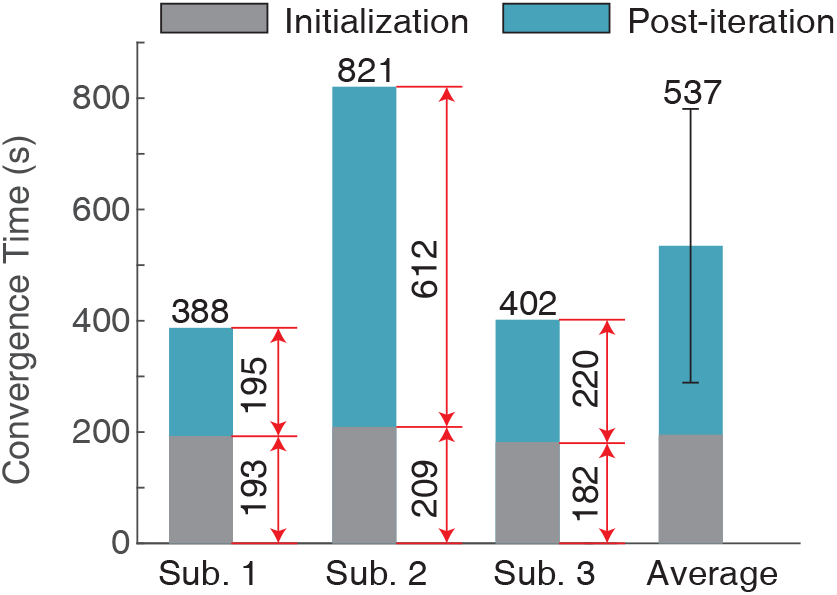
The convergence time represents the time spent in the whole experiment, including the initialization time and the post-iteration time. The initialization time represents the experimental time of the six sets of initialization parameters, and the post-iteration time represents the experimental time of the subsequent iterative process.

For each subject, the six sets of control parameters randomly selected from the six parts evenly divided by the feasible domain of the control parameters are shown in Fig. 8. The initialization parameters of each subject are different. The process of the whole optimization experiment is shown in Fig. 9. The gray shaded parts in Fig. 9 represent the six sets of initialization control parameters during the optimization process, and their corresponding gait symmetry indicator under the action of the six sets of control parameters. The yellow area represents that the gait symmetry indicator value obtained by the iterative process satisfies the convergence condition. For Subject 1, 2 and 3, accordingly, 12, 24 and 13 iterations are needed to reach convergence condition.

**Figure 8.**
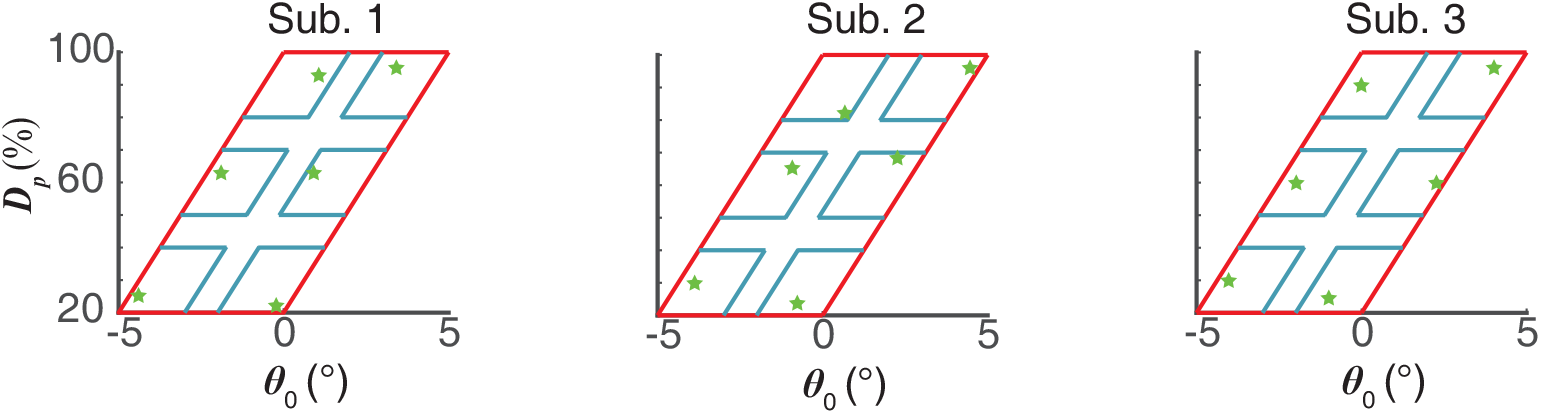
The six initialization control parameters of the subjects. *θ*_0_ is the desired equilibrium angle, i.e. ankle angle when heel strike. *D*_*p*_ determines the relationship between the torque and the speed, it can be regarded as the damping coefficient. The entire feasible area of *θ*_0_ and *D*_*p*_ is divided into six uniform small areas, and six green pentagram represent six sets of initialization control parameters randomly selected in six small areas.

**Figure 9.**
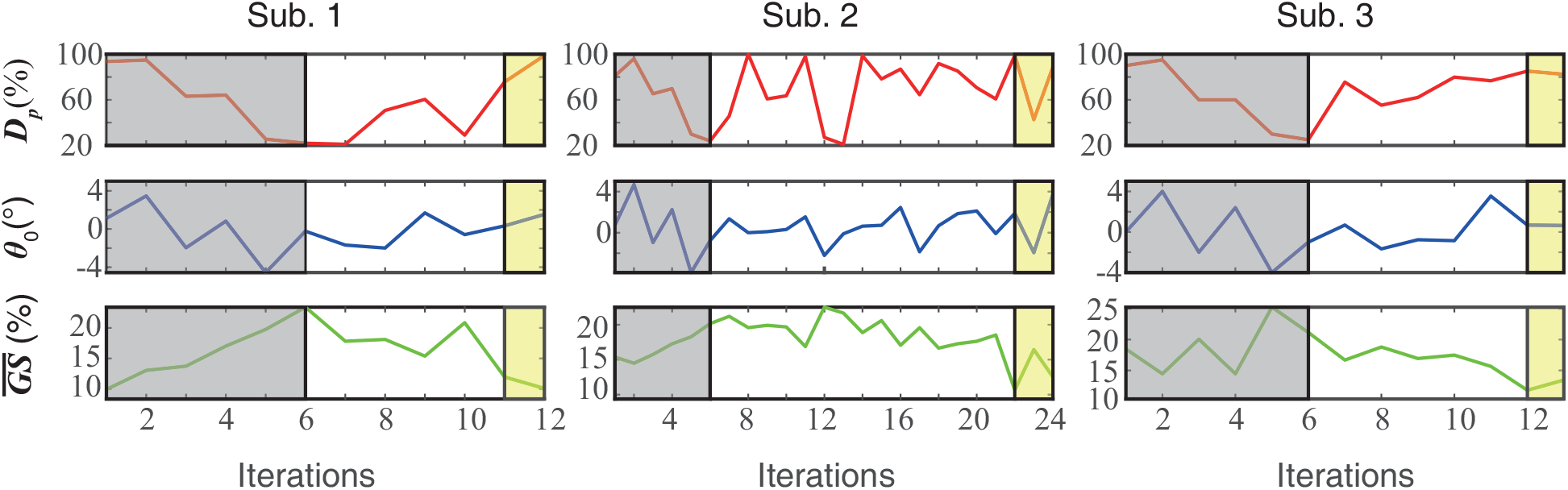
Optimization process. Iterations represent the number of iterative steps in the optimization process, *D*_*p*_ and *θ*_0_ represent control parameters, and 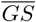 represents the gait symmetry indicator. The gray shaded area represents the initialization process in the optimization process, and the yellow shaded area represents the convergence region of the optimization process.

The gait symmetry indicator and the corresponding control parameters while convergence, show the discrepancy between different individuals. Subject 1 took 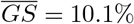 when the control parameters of the robotic prosthesis were *D*_*p*_ = 98.79%, *θ*_0_ = 1.5°, Subject 2 took 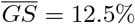 while the control parameters *D*_*p*_ = 88.55%, *θ*_0_ = 3.8°, and Subject 3 took 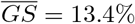 while the control parameters *D*_*p*_ = 82.36%, *θ*_0_ = 0.6°. These results demonstrate individual difference and also the necessity of customizing individual parameters. The maximum value, the intermediate value, and the minimum value of the gait symmetry indicator are utilized to evaluate the optimization performance, in Fig. 9. Also, as the reference, the gait symmetry indicator from the subjects wearing passive prostheses are demonstrated in Fig. 9. Compared with the gait symmetry indicator using the passive prostheses, for Subject 1, 2 and 3, the best gait symmetry indicator was decreased by 52.0%, 12.2% and 6.0% (21.0%, Mean), respectively.

### 7.2 Kinetics

When the subject wears the prosthesis and walks on the treadmill, we measured the GRFs of the left and right feet of the passive prosthesis and the robotic prosthesis in the three sets of control parameters min condition, mid condition, max condition. It can be seen from Fig. 11 that in the max condition, the peak of the GRFs of the intact side is larger than the other three conditions, because the corresponding control parameter *D*_*p*_ is smaller in the max conditon. From Eq. 1, it can be seen that the braking torque value provided by the prosthetic side is small at the condition, so the intact side provides more force, the peak force at the intact side is higher in the max condition.

**Figure 10.**
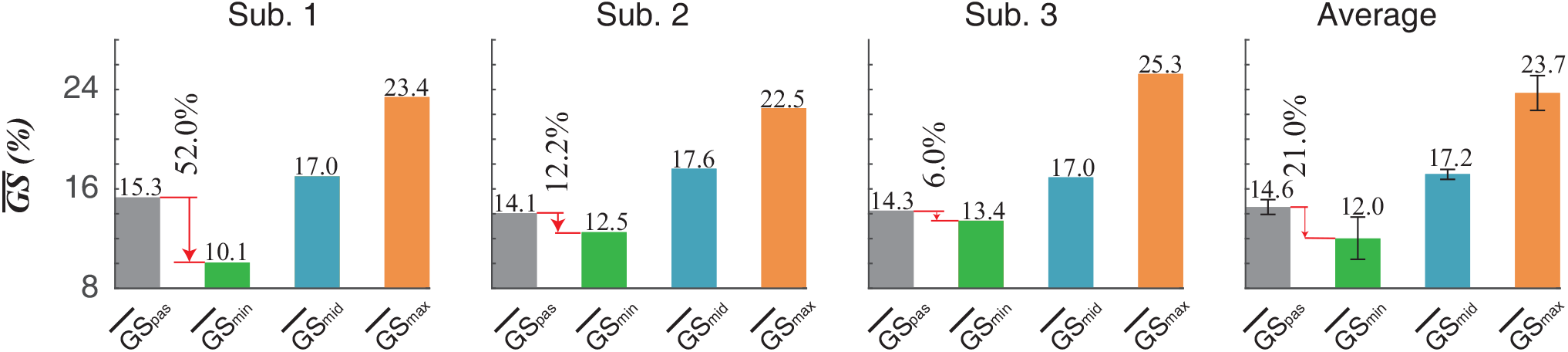
The gait symmetry indicator of the subjects under four conditions. (1) passive represents passive condition, it indicates the condition of a passive prosthesis. (2) min represents min condition, it indicates that the gait symmetry indicator value obtained by the robotic prosthesis under the control parameter is the minimum value in the whole optimization process, that is, the gait symmetry indicator is the best case. (3) mid represents mid condition, it indicates that the gait symmetry indicator value obtained by the robotic prosthesis under the control parameter is the intermediate value in the whole optimization process. (4) max represents max condition, it indicates that the gait symmetry indicator value obtained by the robotic prosthesis under the control parameter is the maximum value in the whole optimization process, that is, the gait symmetry indicator is the worst case. 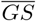 represents the gait symmetry indicator. Average represents the average of the gait symmetry indicator of three subjects

**Figure 11.**
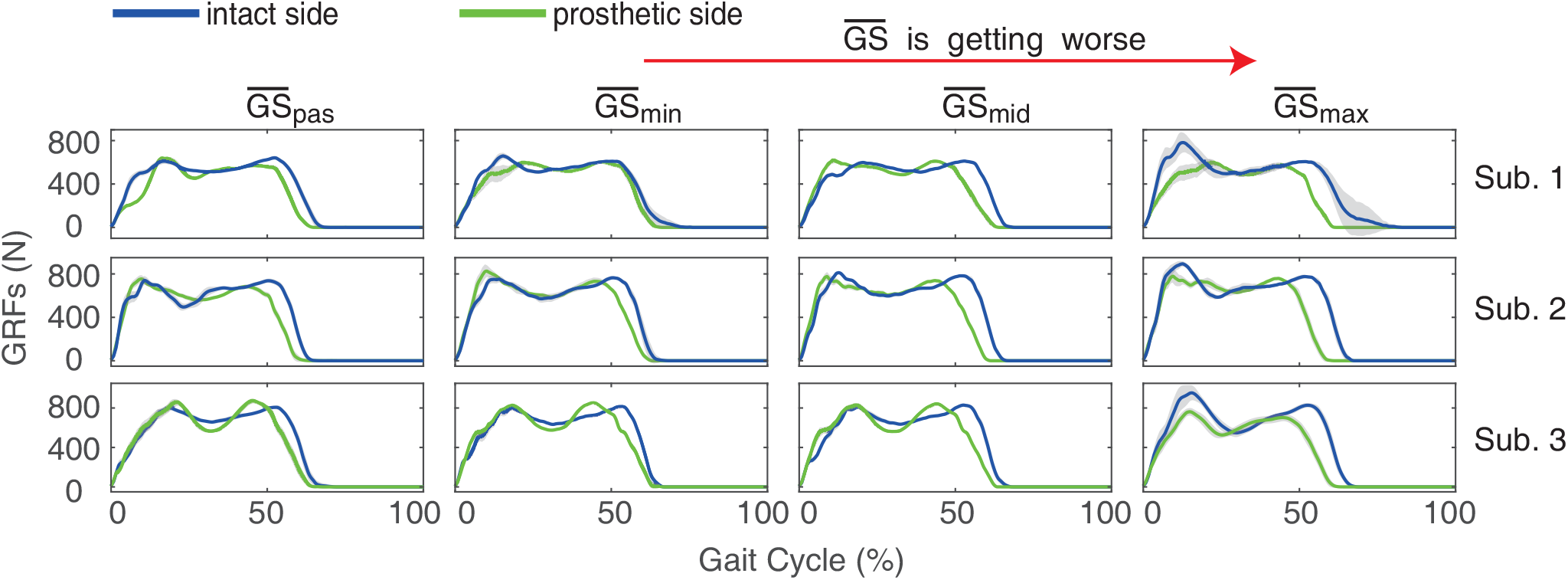
The GRFs of the subjects under four conditions. (1) passive represents passive condition, it indicates the condition of a passive prosthesis. (2) min represents min condition, it indicates that the gait symmetry indicator value obtained by the robotic prosthesis under the control parameter is the minimum value in the whole optimization process, that is, the gait symmetry is the best case. (3)mid represents mid condition, it indicates that the gait symmetry indicator value obtained by the robotic prosthesis under the control parameter is the intermediate value in the whole optimization process. (4) max represents max condition, it indicates that the gait symmetry indicator value obtained by the robotic prosthesis under the control parameter is the maximum value in the whole optimization process, that is, the gait symmetry is the worst case. GRFs represents the ground reaction forces

Moreover, it can be obtained from Fig. 11 that in the min condition, the time ratio of stance phase to swing phase between the intact side and the prosthetic side in one gait cycle is also closer. This also shows that the more similar the dynamic characteristics of the intact side and the prosthetic side, the better the gait symmetry indicator. In the max condition, the prosthetic side has a longer swing period in one gait cycle. Because the robotic prosthesis has a small braking torque when the damping is low, and does not provide sufficient torque, it is passively dragged by the intact side. Therefore, in the max condition, the stance phase of the intact side is longer than the prosthetic side.

When the prosthesis worked under four conditions (i: min condition, ii: mid condition, iii: max condition, iiii: passive condition), the range of ankle angle and knee angle on the prosthetic side of the subject were measured, in Fig. 12. It can be seen from Fig. 12 that in the min condition, the ankle angle is different at the beginning of the gait cycle (heel strike). Because each subject has his uniqueness, the control parameter *θ*_0_ in each of the best control parameters is different, the ankle angle on the prosthetic side is different at heel strike. From Fig. 12 and Tab. 2 in the min, mid and max condition, as the gait symmetry indicator becomes worse, the range of the ankle angle on the prosthetic side increases, because the control parameter *D*_*p*_ is the smallest in the max condition, the range of the ankle angle on the prosthetic side is largest. Moreover, the *SI* value will decrease. It can be seen from Tab. 2 that under the max condition, the *SI*_*max*_ value is the smallest. From the perspective of kinematics, the gait symmetry is the best in the max condition. From Fig. 12 that when the passive prosthesis is worn, the ankle angle of the prosthetic side is not large, but there is still a range of change. Because the carbon-fiber footplate on the passive prosthesis is deformed during walking. For the knee angle range of the subject 2 in Fig. 12, when compared with the passive condition, the knee joint motion characteristics of the intact side and the prosthetic side in the min condition are more symmetrical and have a significant improvement. For subject 1 and subject 3, the knee joint motion symmetry of the knee angle was not significantly improved. This shows that there is no uniform regularity for the improvement of the knee joint motion symmetry of the subjects as the gait symmetry indicator improves. It has individual differences for different subjects.

**Table 2.** Ankle angle and knee angle range on the prosthetic side and intact side.

| Joint range | Subject | Passive |  |  | Min |  |  | Mid |  |  | Max |  |  |
| --- | --- | --- | --- | --- | --- | --- | --- | --- | --- | --- | --- | --- | --- |
| | | Prosthetic (°) | Intact (°) | $SI_{pas}(\%)^1$ | Prosthetic (°) | Intact (°) | $SI_{min}(\%)^1$ | Prosthetic (°) | Intact (°) | $SI_{mid}(\%)^1$ | Prosthetic (°) | Intact (°) | $SI_{max}(\%)^1$ |
| Ankle | 1 | 7.7(0.1) | 28.7(0.5) | 115(2.1) | 16.1(0.5) | 28.5(1.0) | 55.8(4.7) | 18.5(0.7) | 27.3(0.9) | 38.4(2.5) | 26.8(0.7) | 29.3(0.5) | 9.0(3.8) |
|  | 2 | 11.1(0.3) | 32.4(1.0) | 98.2(2.5) | 18.1(0.3) | 36.0(1.6) | 66.2(2.8) | 20.3(0.4) | 30.7(0.5) | 40.7(2.9) | 26.5(0.4) | 36.2(2.9) | 30.9(5.9) |
|  | 3 | 12.2(0.2) | 30.4(0.7) | 85.2(1.1) | 17.2(0.3) | 27.9(0.8) | 47.5(1.7) | 19.9(0.8) | 26.6(1.4) | 28.5(6.5) | 23.0(1.0) | 28.5(1.2) | 21.3(8.3) |
| Knee | 1 | 69.0(1.5) | 67.8(0.6) | 1.7(1.3) | 74.7(4.0) | 62.3(1.0) | 18.0(6.3) | 71.4(1.5) | 58.6(0.4) | 19.6(2.6) | 66.9(1.6) | 60.7(0.9) | 9.8(2.5) |
|  | 2 | 61.1(1.8) | 74.7(1.6) | 20.1(1.4) | 65.3(0.8) | 69.7(0.9) | 6.5(1.6) | 60.6(0.6) | 65.3(0.2) | 7.4(1.3) | 61.9(0.7) | 67.0(0.5) | 7.8(1.1) |
|  | 3 | 59.0(2.9) | 73.7(2.8) | 22.2(7.1) | 58.4(1.0) | 68.9(1.0) | 16.5(1.9) | 57.2(1.0) | 69.4(1.3) | 19.3(2.7) | 57.2(0.9) | 70.8(2.5) | 21.1(4.0) |

<sup>1</sup> $SI = \left| \frac{\theta_P - \theta_I}{\frac{1}{2}(\theta_P + \theta_I)} \right| \times 100\%$ , where $SI$ represents the symmetry index, $\theta_P$ represents the range of ankle or knee of the prosthetic side, and $\theta_I$ represents the range of ankle or knee angle of the intact side Herzog et al. (1989).

**Figure 12.**
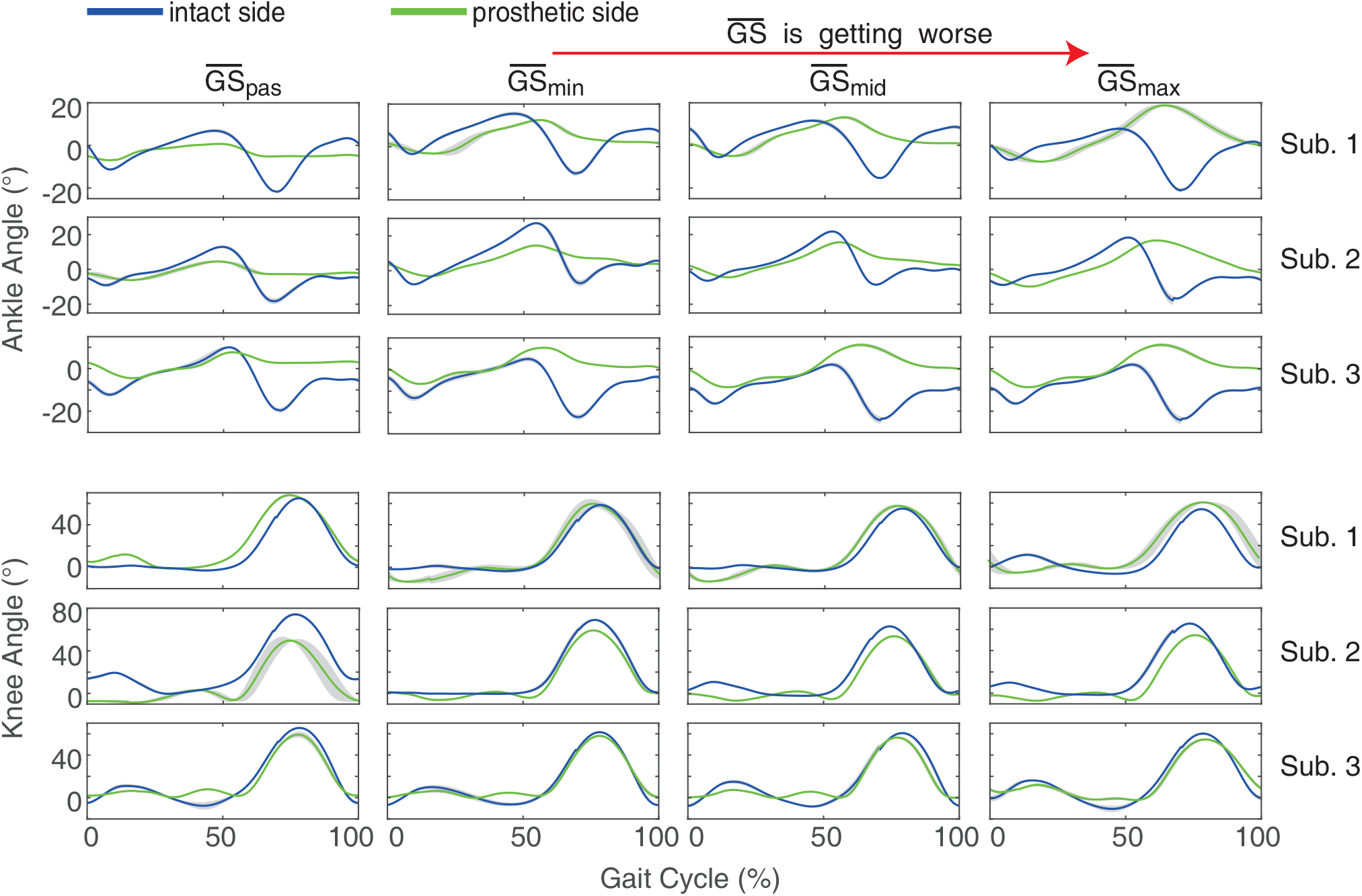
The range of motion of the ankle and knee angles of the three subjects when the prosthesis was operated under four conditions,(1) passive represents passive condition, it indicates the condition of a passive prosthesis. (2) min represents min condition, it indicates that the gait symmetry indicator value obtained by the robotic prosthesis under the control parameter is the minimum value in the whole optimization process, that is, the gait symmetry is the best case. (3)mid represents mid condition, it indicates that the gait symmetry indicator value obtained by the robotic prosthesis under the control parameter is the intermediate value in the whole optimization process. (4) max represents max condition, it indicates that the gait symmetry indicator value obtained by the robotic prosthesis under the control parameter is the maximum value in the whole optimization process, that is, the gait symmetry is the worst case.

### 7.3 Metabolic Cost

Before walking on the treadmill, the subjects stand on the treadmill for at least 2 minutes to measure the static energy consumption of the subjects, and then measure the energy consumption of the subjects during the walking with robotic prosthesis. It subtracts the static energy value to get the net energy consumption *EE*_*net*_ of the subjects. The net energy consumption *EE*_*net*_ obtained by the subjects under passive prosthetic conditions and three sets of control parameters are shown in Fig. 13. From Fig. 13, for the robotic prosthesis, as the gait symmetry indicator of the subject becomes worse, the net energy consumption of the subject increases, except for the mid condition and max condition of the Subject 2, which may be caused by individual differences. This also indicates that the relationship between the gait symmetry indicator and the net energy consumption of the subjects is basically positively correlated. By comparing passive prostheses and robotic prostheses working in the min condition, their net energy consumption is 174.1±14.4*J/*(*kg.min*) and 159.5±8.9*J/*(*kg.min*). Therefore, after optimization, the energy consumption of the wearing robotic prosthesis is less than the net energy consumption of the passive prosthesis. Compared with the net energy consumption using the passive prostheses, for Subject 1, 2 and 3, the best net energy consumption was decreased by 11.0%, 3.0% and 13.4% (9.2%, Mean), respectively. This also shows that although we used the gait symmetry indicator as the optimization target in the experiment, the net energy consumption of the subjects was also optimized.

**Figure 13.**
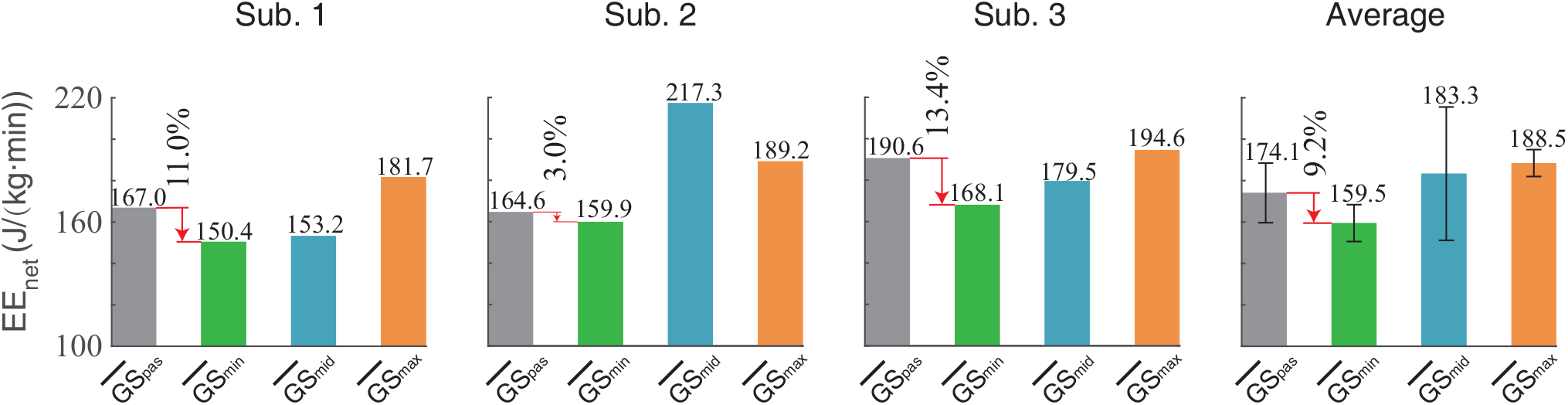
The net metabolic energy consumption value of the subjects under four conditions. (1) passive represents passive condition, it indicates the condition of a passive prosthesis. (2) min represents min condition, it indicates that the gait symmetry indicator value obtained by the robotic prosthesis under the control parameter is the minimum value in the whole optimization process, that is, the gait symmetry is the best case. (3)mid represents mid condition, it indicates that the gait symmetry indicator value obtained by the robotic prosthesis under the control parameter is the intermediate value in the whole optimization process. (4) max represents max condition, it indicates that the gait symmetry indicator value obtained by the robotic prosthesis under the control parameter is the maximum value in the whole optimization process, that is, the gait symmetry is the worst case. *EEnet* represents the net metabolic energy consumption value, Average represents the net metabolic energy consumption value of the gait symmetry indicator of three subjects

## 8. Discussion

### 8.1 Metabolic Cost and Gait Symmetry

This paper investigated the gait-symmetry-based human-in-the-loop optimization, and also the relationship between gait symmetry and metabolic cost was further studied. The majority of results demonstrates that while improving the gait symmetry of the unilateral transtibial amputees wearing the robotic prosthesis, the reduction of the net metabolic energy consumption was accompanied. Only for Subject 2, the net metabolic cost consumption under mid-gait-symmetry condition, is even higher than the net metabolic cost consumption under max-gait-symmetry condition (worst gait symmetry). This result implies that the relationship between gait symmetry and the net metabolic cost consumption is not explicit or individual traits may affect this relationship. In spite of these implicit relationships between gait symmetry and the net metabolic cost consumption, the tendency is clear that the net metabolic cost consumption under the best-gait-symmetry condition is far lower than the consumption under the worst-gait-symmetry condition.

For the prostheses, too heavy weight may result in the gait asymmetry and increase the metabolic cost Smith and Martin (2013). In this paper, the robotic prosthesis (1.58*kg*) is heavier than the passive prosthesis (0.408*kg*). Although the prosthesis only provides the braking torque (about 50*Nm* Yuan et al. (2017)) and cannot provide the enough push-off torque (the able-bodied: 120*Nm* Winter (1991)). But this paper demonstrated that after optimization, compared with the walking performance using the lighter-weight passive, after optimization, using the robotic prosthesis, the gait symmetry was improved by 21.0% and the metabolic cost was reduced by 9.2%. This result demonstrates that the robotic prosthesis without the push-off power has the potential ability to improve walking performance in gait symmetry and metabolic cost.

### 8.2 Gait Symmetry Based on Kinetics and Kinematics

There are many gait symmetry indicators to evaluate the walking performance in some aspacts Kendell et al. (2010). In this paper, we use the root mean square value of amputees’ GRFs as the gait symmetry indicator. The gait symmetry indicator is used as the objective function in the optimization process, and finally the optimal control parameters of the robotic prostheses are obtained. Using this optimal control parameter, the net metabolic energy consumption can be reduced compared to passive prostheses. The angle of the ankle joint on the prosthetic side of the amputee can also be used as a gait symmetry indicator to measure the amputees’ effect during the walking process. The larger the range of the ankle joint angle of the prosthetic side, the better the gait symmetry indicator of the amputee. It can be seen in Fig. 12 that under the three sets of control parameters min, mid, max, the measured prosthetic side ankle joint angle is sequentially increased. Therefore, if the ankle joint angle range of the amputee’s prosthetic side is used as the gait symmetry indicator, the gait symmetry indicator is gradually improved. This is exactly the opposite of the result obtained by using the root mean square value of GRFs as the gait symmetry indicator. At the same time, studies have shown that when the kinematic parameters are used as the gait symmetry indicator of the amputee and the kinetic parameters are used as the gait symmetry indicator, the results are opposite Childers and Kogler (2014). Combined with the measured net metabolic energy consumption value analysis, the net metabolic energy consumption value at this time is reduced. Therefore, from the perspective of reducing the metabolic energy consumption of the amputee, the root mean square value of amputees’ GRFs is used as the gait symmetry indicator. It is more conducive to reducing the metabolic energy consumption value. This may also mean that the root mean square value of amputees’ GRFs is more closely related to their own metabolic energy consumption than the range of the prosthetic side ankle angle. From the perspective of reducing metabolic energy consumption, the use of force-related gait symmetry indicators may be more effective.

### 8.3 Convergence time

In the gait-symmetry-based human-in-the-loop experiment, which uses the gait symmetry indicator of the subject as the optimization target, the convergence time of the entire optimization experiment of all the subjects 9.0 4.1*min*. The optimization experiment of human-in-the-loop with energy consumption as the optimization goal has an average convergence time of 21.4±1.0*min* Ding et al. (2018). This may be because when we measure the gait symmetry indicator, we only need 25 strides to calculate a gait symmetry indicator, and the time for each stride of the subject is relatively short, so the time spent on each iteration is about 30s. For the metabolic energy measurement of the subjects, it is necessary to measure the stable metabolic energy consumption value of the subject during walking, which takes an average of 2 min Felt et al. (2015), This may be the reason why the convergence time of the experiment with the gait symmetry indicator as the optimization target is relatively short. The shorter convergence time helps us to shorten the experiment time and reduce the impact of the fatigue of the subject during the experiment process.

### 8.4 Limitation

In our prosthetic control process, compared with the able-bodied (120*Nm* Yuan et al. (2017)), the prosthetic limb cannot provide enough push-off torque (only about 50*Nm* Yuan et al. (2017)) during stance phase, which may also affect the results of the optimal control parameters obtained during our optimization process. The robotic prosthesis used in our experiment doesn’t have the push-off control stage, it may also affect the amputees’ gait symmetry and energy consumption. The robotic prosthetics with push-off control stragey can better improve the energy consumption of the amputees Feng and Wang (2017), and also make the ground reaction forces of the amputees different while walking. Therefore, whether the best control parameters of the robotic prosthesis can be obtained still needs further consideration. If the feasible domain of the control parameter may be expanded, the convergence time may be prolonged and the corresponding control parameters will be altered. More importantly, taking the safety into consideration, the feasible domain of the *D*_*p*_ and *θ*_0_ control parameters is now limited as a parallelogram region. Meanwhile, the corresponding risk of injury will also increase, *e.g*. due to insufficient damping braking, the fall off.

## 9. Conclusion

In this paper, we proposed gait-symmetry-based human-in-the-loop optimization. Three unilateral transtibial subjects participated in the experiments. Compared to gait symmetry while wearing prosthetics, after optimization, the gait symmetry indicator value of the subjects wearing the robotic prostheses was improved by 21.0% and meanwhile the net metabolic energy consumption value was reduced by 9.2%. The experimental results show that the convergence time takes about 9.0±4.1*min*. Compared with the metabolic energy consumption as the optimization target (one iteration: 2*mins*), the period of each iteration while using the gait symmetry as the optimization target could be shorten to 30*s*. From a biological point of view, the better the gait symmetry indicator of walking with robotic prostheses, the more uniform the force on the legs and the less damage to the prosthetic side of the amputee. It may be more conducive to the recovery of the amputees. Therefore, in the optimization experiment of robotic prosthesis, it may be more feasible and efficient to adopt gait symmetry indicator as the optimization goal. Further, the experimental results show that the improvement kinetics-based gait symmetry and the reduction of metabolic cost consumption have a certain positive correlation. In contrast, the improvement kinematics-based gait symmetry and the reduction of metabolic cost consumption have a certain negative correlation. The positive/negative correlation indicates that choosing gait symmetry calculated by kinetic characteristics may be more reasonable. In the future, the gait-symmetry-based human-in-the-loop optimization method paves the way for online optimization and may have a wider application.

## Acknowledgement

This work was supported by the National Key R&D Program of China (No. 2018YFB1307302) and the National Natural Science Foundation of China (No. 91648207, No. 51922015, No. 91948302).

